# Foraging gene and xenobiotic stresses relationships in *Drosophila melanogaster*

**DOI:** 10.1101/2020.07.24.219550

**Authors:** Amichot Marcel, Tarès Sophie

**Affiliations:** Université Côte d’Azur, INRAE, CNRS, ISA, France

**Keywords:** Foraging, cGMP-dependent protein kinase, xenobiotic, adaptation, insecticide, drosophila

## Abstract

In several insect species, the foraging behaviour has been demonstrated to be controlled by the *foraging* gene (*for*) which encodes for a cGMP-dependent protein kinase (PKG). In wild Drosophila populations, rover and sitter individuals coexist and are characterized by high and low PKG activity levels respectively. Because of their increased foraging behaviour, we postulated that rover flies are more exposed to environmental stresses than sitter flies. Thus, we tested whether rover and sitter flies differ in their fitness by using insecticides as chemical stressors. We showed that their responses are different depending on the insecticide used and are linked to variations of cytochrome P450s activities. Furthermore, the expression of the insecticide metabolizing cytochrome P450 *Cyp6a2* was shown to be under the control of the *for* gene. We evidence here a new physiological function for the *for* gene in Drosophila and we demonstrate its involvement in the adaptation to chemicals in the environment.

## Introduction

Foraging is a vital behaviour for animals to ensure their survival. However, this behaviour is the result of a trade-off between benefits, expenses and risks associated with foraging. This has been formalized in the Optimal Foraging Theory which is based on several founder papers (Charnov, 1976; Emlen, 1966; Fretwell & Lucas, 1970; MacArthur & Pianka, 1966; Schoener, 1971). We put forth the hypothesis that foraging can also lead to an enhanced exposure to compounds present in the environment.

Searches for genes that control the foraging behaviour have been successful in Drosophila. One series of strains was remarkable as a single locus was demonstrated to control foraging to define rover and sitter populations (Pereira & Sokolowski, 1993). Rover larvae continue to explore the environment even if they have found food whereas sitter ones stop their exploration when on food (Kaun & Sokolowski, 2009). A single gene, *for*, was associated with this locus. It encodes for a cGMP-dependent protein kinase (PKG) and its over expression demonstrated that it was responsible for the foraging behaviour (Osborne et al., 1997). Widely distributed in eukaryotes, PKG has been described to affect various functions in the brain or in the cardiovascular system (Hofmann, 2006). In Drosophila, the PKG is involved in memory (Mery et al., 2007), heart contractions (Johnson et al., 2002), axon guidance (Peng et al., 2016) or nociception escape (Dason et al., 2019). In wild Drosophila populations, the foraging behaviour is naturally polymorphic with a quite constant proportion of rover and sitter flies (70% rovers, 30% sitters) (Sokolowski, 1980) although this may vary depending on the population density ((Sokolowski et al., 1997) or the allele frequency (Fitzpatrick et al., 2007). Furthermore, the *for* locus also control the dispersal of adults (Edelsparre et al., 2014). Thus, the *for* gene is involved in numerous and various physiological and behavioural processes.

While foraging and feeding, insects have to manage with variable chemical stresses. Insects have been successfully faced with allelochemical compounds throughout the evolution mainly by means of specialized enzymes involved in the excretion or the degradation of these compounds. To all these naturally occurring stresses, humans have added xenobiotics and especially insecticides. Their recent use has evidenced the swiftness and the efficiency of the adaptation ability of insects. Indeed, insects challenge our chemical arsenal by several molecular means including insecticide target modification, increased excretion or enhanced metabolism of the insecticides resulting into resistances (Dawkar et al., 2013). The metabolism of allelochemical compounds and xenobiotics is mediated by enzymes belonging to three categories: esterases (EST), glutathion transferases (GST) and cytochrome P450s (P450). As far as we know, each category is present in every insect species so that an insect can have dozens of P450s, GSTs and esterases (Feyereisen, 2005). Some of the enzymes active against allelochemicals are also able to metabolize xenobiotics (for instance, see (Li et al., 2007)).

In this study, we tested whether adaptation abilities to chemical stresses are different between rover and sitter flies. The availability of mutants and transgenic flies with modified foraging behaviour makes Drosophila a very suitable model for this study. The toxicological and metabolic fates of the insecticides we used here as environmental stressors are also well known in Drosophila. Our results demonstrate that the *foraging* gene modulates the adaptation ability prior to any exposure to environmental chemicals.

## Material and methods

### Drosophila strains

Two rover strains (*for^R^, dg2*-cDNA) and three sitter strains (*for^S2^, w^1^;for^S^* and *for^S^*) were provided by Prof. MB. Sokolowski (University of Toronto, Canada). The *for^R^* and *for^S^* strains were selected from a single wild population by sorting rover and sitter larvae according to their foraging behaviour (Pereira & Sokolowski, 1993). After applying a mutagenic protocol on *for^R^* flies, a selection for a sitter phenotype was performed in the progeny to establish the *for^S2^* strain. It was demonstrated that the *for^R^* and *for^S2^* strains differ only by the *for* gene (de-Belle et al., 1989; Pereira & Sokolowski, 1993) so that *for^S2^* is hypomorphic regarding the *for* gene. The rover *dg2*-cDNA strain was derived from the sitter *w^1^;for^S^* strain after transgenesis using a *for* cDNA (T2 transcript) which is under the control of a leaky heat-shock protein promoter. So, the *dg2*-cDNA strain is a gain-of-function strain compared with *w^1^;for^S^* (Osborne et al., 1997). Because of this leakiness and the fact that temperature interferes with toxicological measures (Gordon et al., 2014), we chose to avoid the heat shock treatment. We thus had at our disposal two pairs of genetically tightly linked strains, *for^R^* and *for^S2^* on one hand and *dg2-cDNA* and *w^1^;for^S^* on the other hand, differing only by the foraging behaviour associated with the expression level of the PKG. For this reason, we decided to consider only the *for^R^/for^S2^* and *dg2*-cDNA/*w^1^;for^S^* pairs of strains. Nevertheless, all the results obtained with the *forS* strain are given as supplementary material.

All flies were reared on standard corn-meal medium, 20°C, 70% humidity, 12/12 h photoperiod. For every experiment, the same number of males and females were used.

### Toxicological tests

To test the adaptation ability of the Drosophila strains, we selected three insecticides: aldrin, deltamethrin and diazinon. These insecticides belong to different chemical families and have various targets in the nervous system. Indeed, aldrin is an organochlorine insecticide and is active on the GABA receptor, deltamethrin is a pyrethroid and is active on the voltage-dependent sodium channel and diazinon is an organophosphorous compound active on the acetylcholine esterase enzyme (Casida, 2009). Moreover, the metabolic fates of these insecticides and their toxicological consequences are well known in insects (Gilbert, 2012; Pfafflin & Ziegler, 2012; Pisani-Borg et al., 1996; Soto & Deichmann, 1967; Stevenson et al., 2011). Indeed, these insecticides can be metabolized by ESTs, P450s and GSTs. EST enzymes can cut ester bounds. P450s add an oxygen atom to the substrate which then undergoes self-rearrangement to give the final product(s) of the reaction. GSTs catalyse the nucleophilic conjugation of the glutathion with the molecules to make them excretable. All these reactions generally give less toxic compounds excepted for the aldrin as its metabolite (dieldrin) is more toxic than the starting molecule (Figure 1).

**Figure 1.**
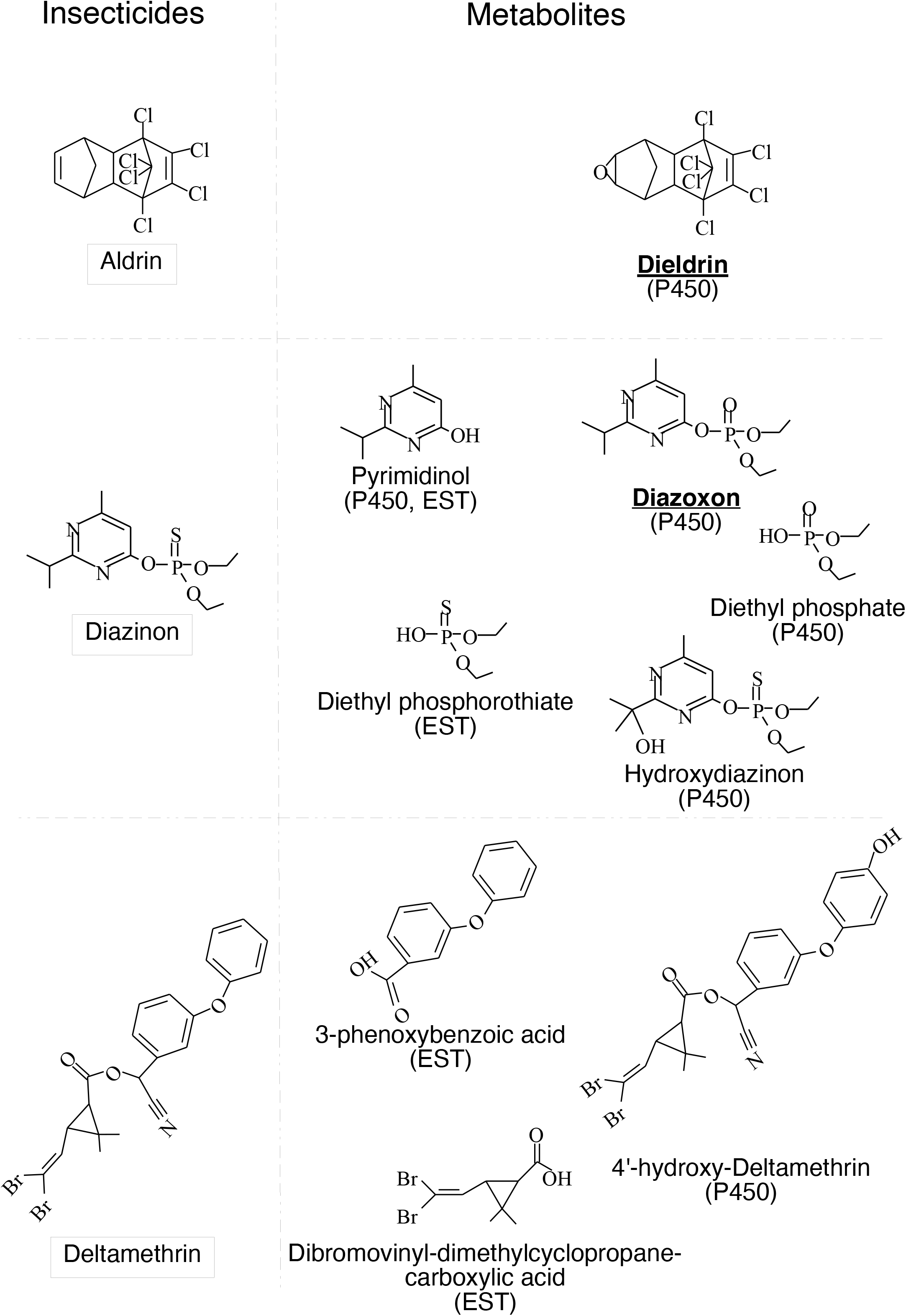
Main EST and P450 metabolites of Aldrin, Diazinon and Deltamethrine in insects. The structural formulas of the three insecticides are presented together with those of the most frequent metabolites observed after the action of EST or P450. The names of the toxic metabolites are bold and underlined. For the clarity of the figure, the GST metabolites were not presented.

The tarsal contact technique was used to test the response of the Drosophila strains to the insecticides. Glass tubes (20 cm2) were coated with 50 *μ*L of an acetonic solution of the tested insecticide at the selected concentration and allowed to dry under a fume hood for 2 hours. Control tubes were treated only with acetone. Ten fies were placed in each tube, both males and females. These tests were repeated three times adding up from 90 to 140 flies for each insecticide concentration and strain. For each insecticide, five concentrations were tested. Drosophila mortality was scored after a 4 h period. We used the Priprobit software (Priprobit 1.63 © Masayuki Sakuma) to calculate and statistically compare the LC50s (concentration that kills half of the population expressed as the insecticide concentration in the solutions used to treat the tubes)) for each strain. When tested, the cytochrome P450 inhibitor piperonyl butoxide (1 mM final) (PBO)(Hodgson & Levi, 1999) was incorporated in 1 mL of corn-meal medium placed in the glass tube already coated with the insecticide solution.

### Enzymatic activities measurements

ESTs, GSTs and P450s activities were tested with substrates that are not specific for a unique enzyme within a family but are able to evidence global variations in the activity of the family. All the protocols, adapted for working with a single fly or with one abdomen in micro plates wells, were based on our experience or on previously published protocols (de Sousa et al., 1995; Nauen & Stumpf, 2002; Pasteur et al., 1997; Zhou et al., 2003).

For all the enzymatic activities measurements, the flies were first anaesthetized with CO_2_, sorted according to their gender and then decapitated to remove the ocular pigments. The same number of males and females were sorted to avoid any sexual dimorphism bias. To determine the baseline values for each enzymatic measure, ten flies were placed in the relevant reaction buffer (see below) and incubated at 100°C for 10 minutes. Then, these flies were processed as described below for each enzyme family. The baseline values are subtracted from the relevant measures to get the activities values. For all of the measurements, we used 96-wells microplates with a single fly or abdomen per well and the activities were expressed as arbitrary units.

To measure esterases activities, each fly was crushed in 180 *μ*L of HEPES (50 mM; pH 7.0) supplemented with 1 *μ*L of a bovine serum albumin (1 mg/mL). Twenty *μ*L of substrate solution were added (alpha- or beta-naphthylacetate 1 mM in water/ethanol (99/1, vol./vol.)). Incubation took place at 30°C for 30 min.; 20 *μ*L of a fast garnet/SDS solution (10 mM each) were added to stop the reaction. The optical density reading of the plate was done with a Spectramax plus384 plate reader (Molecular Devices) at 550 or 490 nm for alpha- or beta-naphthylacetate derivatives respectively.

GSTs activities were measured as follows: each fly was crushed in 50 *μ*L of HEPES buffer (50 mM; pH 7.0). One hundred and fifty *μ*L of glutathione (4 mM in HEPES 50 mM; pH 7.0) and 2 *μ*L of monochlorobimane (30 mM in DMSO) were added. Control incubations were done either with heated flies or without fly or without GSH. Fluorescence (excitation: 390 nm; emission: 465 nm) was measured every 5 min. for 40 min with a Cary Eclipse fluorescence spectrophotometer (Varian). We used at least 40 flies per strain in three different tests.

P450s activities were measured using 7-Ethoxy-Coumarin-O-Deethylase (ECOD). Freshly cut abdomens were individually placed in a well containing 50 *μ*L of phosphate buffer (50 mM; pH 8.0) and 1 *μ*L of a bovine serum albumin (1 mg/mL). The reaction was started by adding 50 *μ*L of phosphate buffer and 1 *μ*L of ethoxycoumarine (20 mM in methanol). There were three kinds of controls: wells without abdomens, wells containing one abdomen supplemented with piperonyl butoxide (5 *μ*L of a 100 mM solution) and wells containing heated abdomens. Fluorescence was measured after a 2 hours incubation at 30 °C and the addition of 100 *μ*L of a v/v mixture of glycine 100 μM pH=10.4 and ethanol (excitation: 390 nm, emission: 450 nm). Tests were repeated 3 times and measures were performed using 24 flies per strain.

### Statistical analysis of the enzymatic activities measurements

We compared the enzymatic activities levels pairwise with the Wilcoxon test using the R software (www.R-project.org). We chose this test because of the number of flies tested (at least 25 per activity) and because it was senseless to test all the strains together because of their genetic history.

In addition, in the supplementary data, we present a comparison of the enzymatic activities levels by visualizing the strain effect size for each activity (Gardner & Altman, 1986) using the DABEST (data analysis with bootstrap-coupled estimation) web application (available at https://www.estimationstats.com, (Ho et al., 2019)). This figure also includes the values measured with the *forS* strain.

### Northern blotting

Total RNAs were extracted using Trizol® (Invitrogene) according to the manufacturer’s protocol. Electrophoretic RNA separation, visualization and transfer was performed according to (http://www.protocol-online.org/cgi-bin/prot/view_cache.cgi?ID=787). The probes were labelled using the PCR DIG probe synthesis kit from Roche. Sequences of the oligonucleotides used to synthesize the probes are listed in the Table 1. Northern blot analyses were performed according to the specifications from Roche. Hybridization and signal detection on the blots were done according to the high stringency conditions described in the protocol supplied by Roche.

**Table 1:**
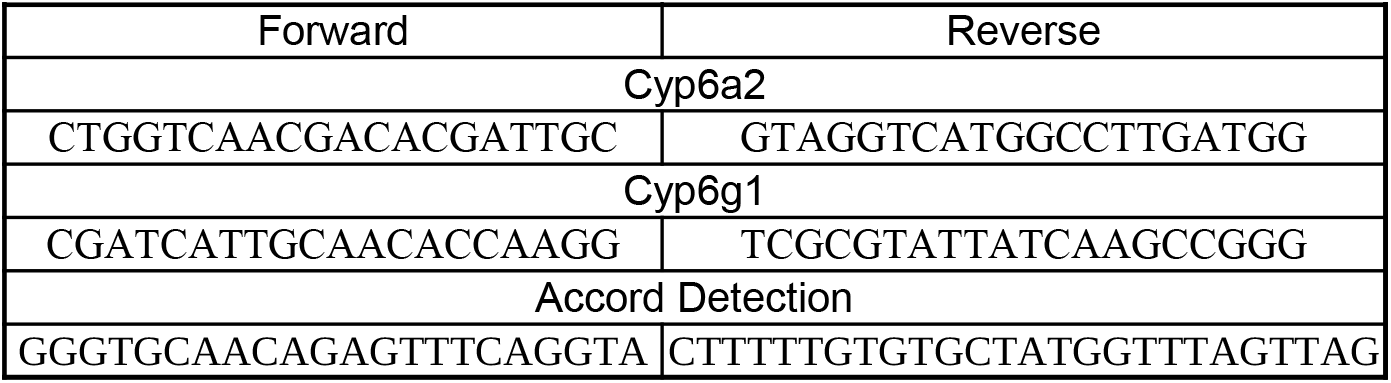
PCR primers sequences The sequences of the *Cyp6a2* and *Cyp6g1* primers were designed using the AmplifX software (Julien N.). The Accord primers were previously published by Daborn et al (2002)

### Accord insertion detection

DNAs were extracted using the VDRC protocol available at (https://stockcenter.vdrc.at/images/downloads/GoodQualityGenomicDNA.pdf). The PCRs to detect the accord insertion in the *Cyp6g1* promotor were performed as described in (Daborn et al., 2002), the sequences of the primers are given in the Table 1.

## Results

### Foraging and the response to chemical stresses in *Drosophila melanogaster*

To test whether the *foraging* gene could modify the response to chemical stresses, we calculated the LC50 of the rover and sitter strains for three insecticides belonging to three different chemical classes and having different targets (Figure 2).

**Figure 2.**
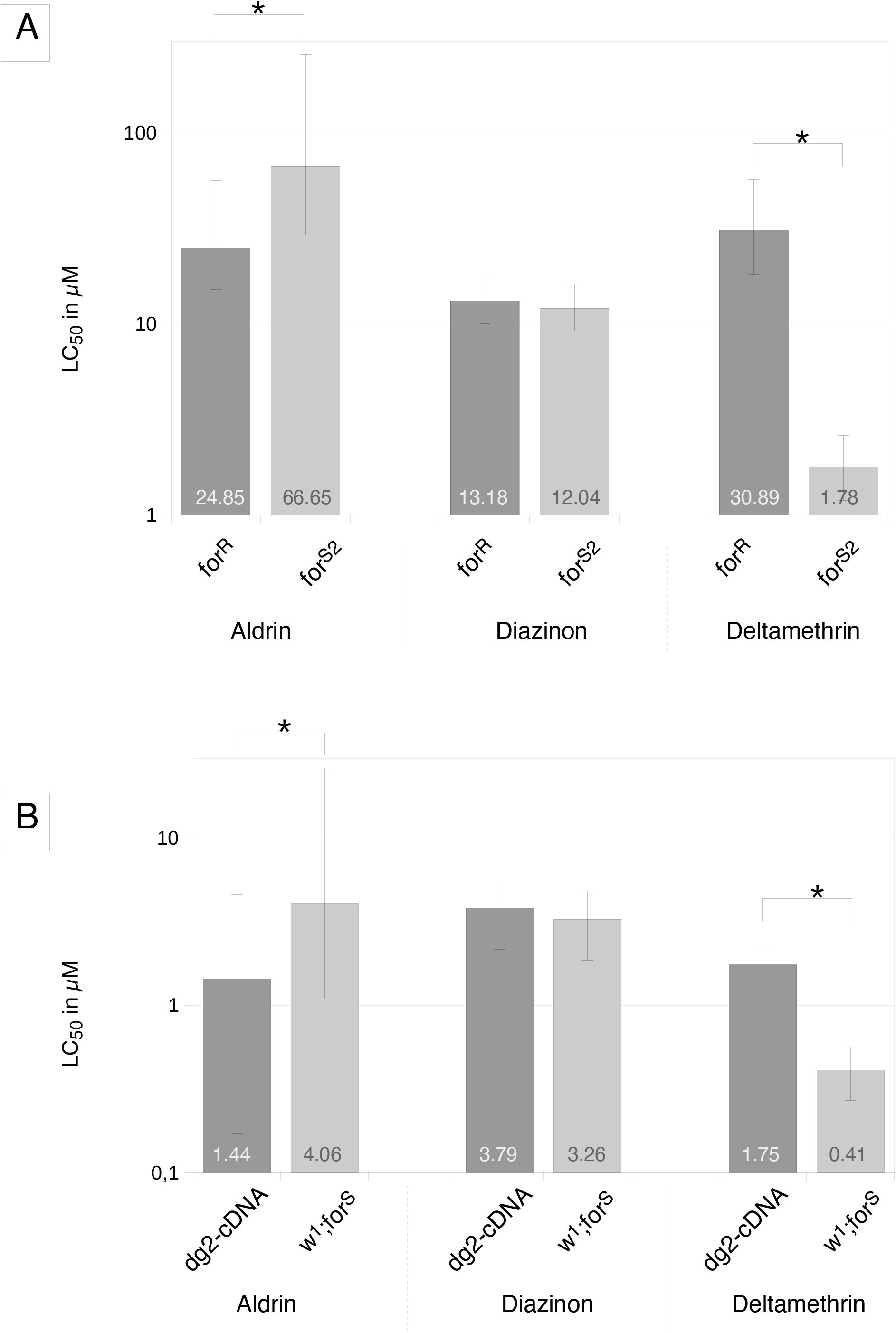
LC_50_ values of the rover and sitter strains for three insecticides Bar charts represent the LC_50_ values of *for^R^* and *for^S2^* strains (panel A) and *dg2-* cDNA and *w^1^;for^S^* strains (panel B). The LC_50_ values are given inside or above the boxes and are expressed in the same units as the insecticide concentrations of the acetonic solutions used to coat the tubes. The vertical bars represent the 95% fiducial limits of the LC_50_ values. The dark grey boxes indicate the rover strains and the light grey boxes the sitter ones. The star indicate significantly different LC_50_ values.

With the organochlorine aldrin, we observed that the rover flies were more susceptible than the sitter ones in both pairs of strains. We found the qualitatively opposite result with the pyrethroid deltamethrin, the rover strains being more resistant than their sitter counterparts. With the third insecticide, the organophosphorous diazinon, we were not able to differentiate between the rover strains and the sitter ones whatever the pair. Thus, the responses of the strains to the insecticides are in relation to their behavioural status although no general rule could be defined yet.

### Xenobiotics degradative enzymes

We wanted to test whether the toxicological differences we observed between the strains could be related to any variation in one of the enzymatic activities known to be responsible for insecticides metabolization (Hemingway & Ranson, 2000). Among the activities we tested, alpha- or beta-esterases and glutathione-S-transferases activities did not vary significantly between the rover and sitter flies within each pair of strains (Figure 3).

**Figure 3.**
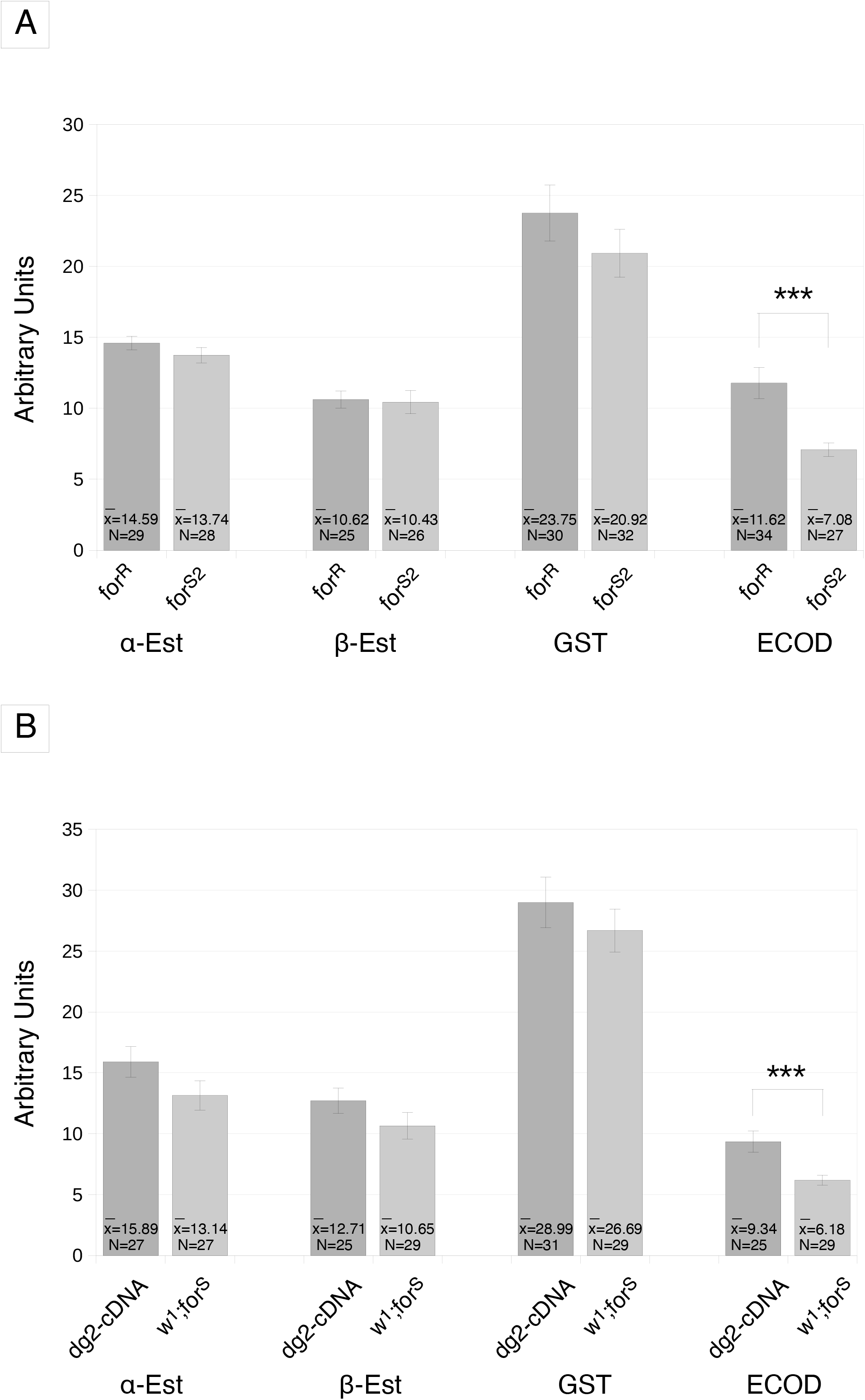
Esterases, GST and ECOD activity levels. Esterases activities are expressed as O.D. arbitrary units/fly/30 min, GST activity is expressed as fluorescence arbitrary units/abdomen/40 min and ECOD activity is expressed as fluorescence arbitrary units/abdomen/120 min. The results for genetically related strains are gathered in panel A (*for^R^* and *for^S2^*) or in panel B (*dg2-cD*NA and *w^1^;for^S^*). The vertical bars represent the standard error of the measures. The three stars indicate the activity values significantly different at p<0,005. The dark grey boxes indicate the rover strains and the light grey boxes the sitter ones.

By contrast, the *for^R^* and *dg2-cDNA* flies, i.e. the rover ones, had a significantly higher ECOD activity compared withshows that the expression of *for^S2^* and *w^1^;for^S^* respectively, i.e. their sitter counterparts (Figure 3). So, this activity linked to P450s is a feature related to the flies behaviour.

### Cytochrome P450s activities, foraging and toxicology

We performed toxicological tests in the presence of a cytochrome P450s inhibitor, PBO, to check whether the P450s are indeed involved in the toxicological differences we observed between the rover and sitter flies. To do such, we fed *for^R^* and *for^S2^* flies, the most different rover and sitter flies regarding toxicology and ECOD activity, with PBO and performed toxicological tests with aldrin, deltamethrin and diazinon (Figure 4). This resulted in the disappearance of the toxicological differences we previously recorded as the new LC50s are now similar between *for^R^* and *for^S2^* in the presence of PBO whatever the insecticide. We evidence here that cytochrome P450s are directly involved in the resistance levels differences observed between these two strains. Thus, we link the toxicological features, the P450 activity levels and the foraging behaviour in these drosophila strains.

**Figure 4.**
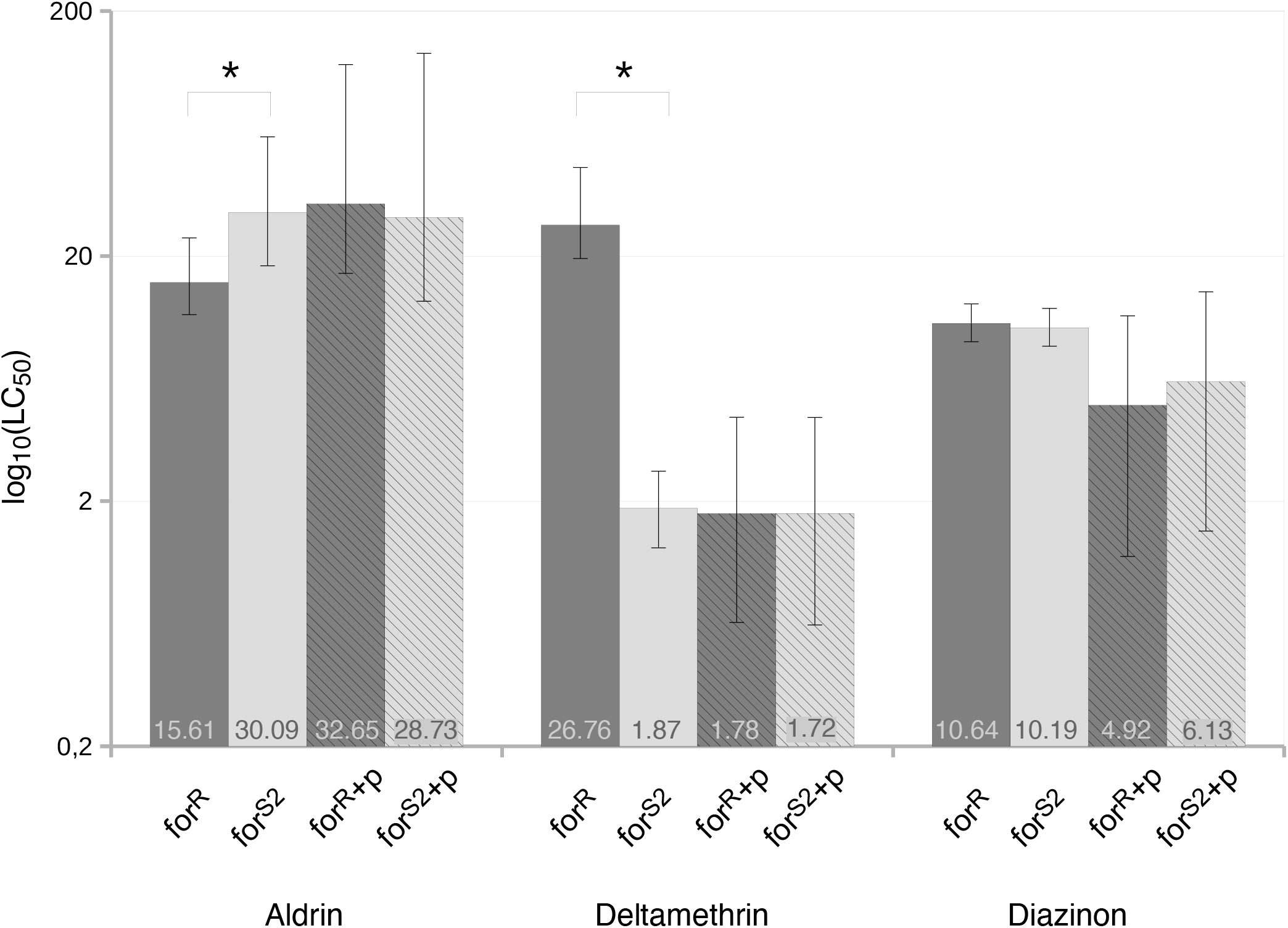
Cytochrome P450 inhibition and tolerance to insecticides The bar chart represents the effect of piperonyl butoxide on the LC_50_ of the *for^R^* and *for^S2^* strains for three insecticides. The LC_50_ of the strains are given in the boxes. Hatched boxes indicate that the flies were treated with piperonyl butoxide. The vertical bars represent the 95% fiducial limits of the LC_50_ values. The dark grey boxes indicate the rover strains and the light grey boxes the sitter ones. The star indicate significantly different LC_50_ values.

**Figure 5.**
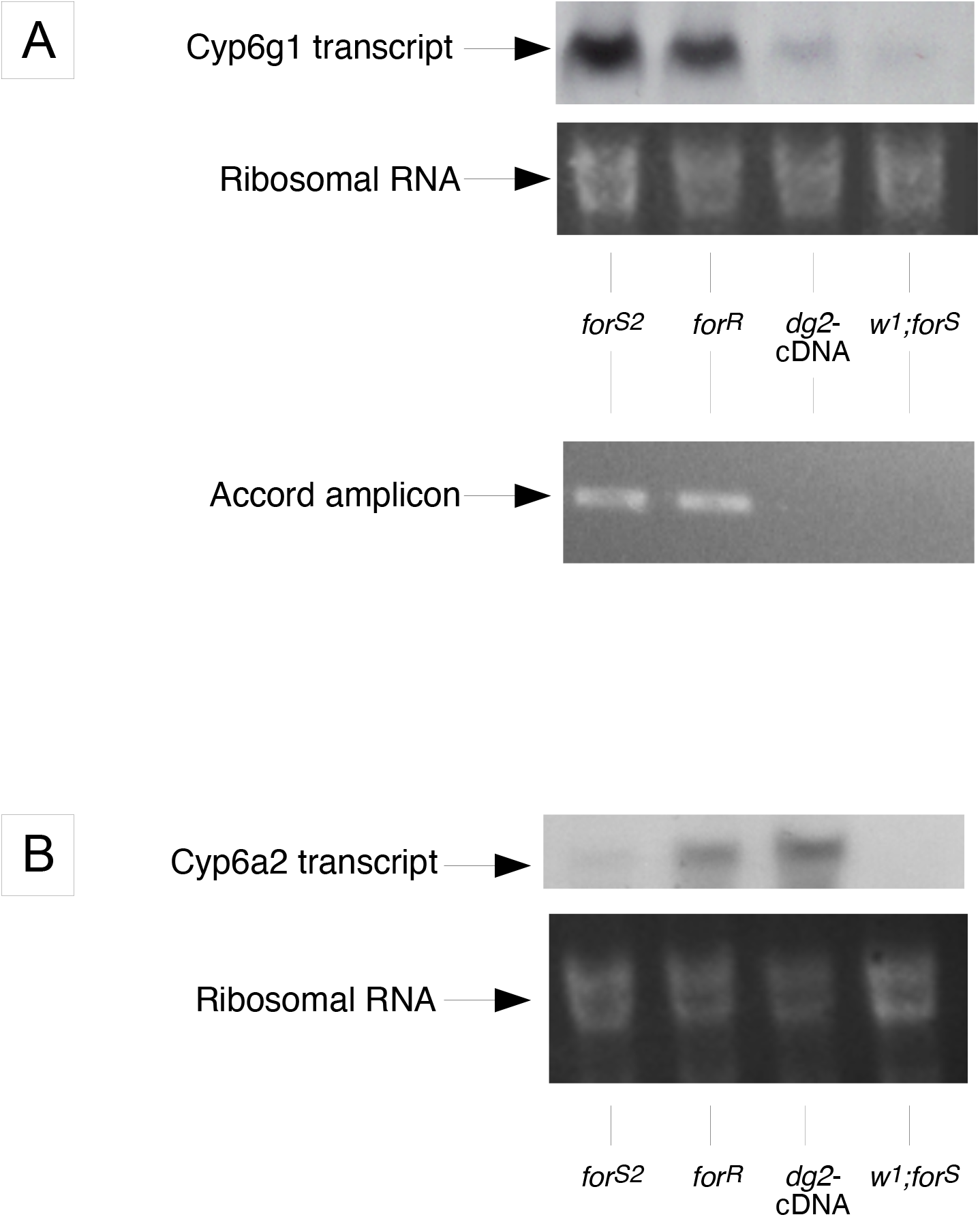
Expression profiles of *Cyp6g1* and *Cyp6a2* (A) Northern blot detection of the *Cyp6g1* transcript and PCR tracking of the *Accord* insertion in the *Cyp6g1* promoter. (B) Northern blot detection of the *Cyp6a2* transcripts. In both panels, the 18S RNA insert demonstrates a similar RNA load in the lanes.

### Cytochrome P450 genes expression

In Drosophila, several cytochrome P450 genes have been linked to insecticide resistance but we focussed our work on two P450s highlighted as insecticides metabolizers: *Cyp6a2* and *Cyp6g1* (Dunkov et al., 1997; Joußen et al., 2010). We compared their expression by Northern blot in the rover and sitter strains.

The figure 4A shows that the expression of *Cyp6g1* is important but similar in *for^R^* and *forS2.* We tested by PCR the presence of an *Accord* element insertion in the *Cyp6g1* promoter sequence as this was previously demonstrated to be responsible for the over expression of this cytochrome P450 (Chung et al., 2007). The *for^R^* and *for^S2^* strains are positive for this test (figure 4A). By contrast, there are neither detectable expression of *Cyp6g1* nor *Accord* element insertion in the *dg2*-cDNA and *w^1^;for^S^* strains.

The situation is different for *Cyp6a2* as it is clearly expressed at a higher level in the two rover strains *for^R^* and *dg2*-cDNA than in their sitter counterparts *for^S2^* and *w^1^;for^S^* (fgure 4B). Thus, the P450 *Cyp6a2* expression level is also linked to the foraging behaviour.

## Discussion

Toxicological analysis of the rover and sitter strains was the first step to verify our working hypothesis. When considering the genetically related rover and sitter strains, i.e. *for^R^* and *for^S2^* on one hand and *dg2*-cDNA and *w^1^;for^S^* on the other hand, we concluded that rover flies were more susceptible to aldrin but more resistant to deltamethrin whereas there was no difference between strains for diazinon. So the effect of the *for* gene seems to be complex because it can be positive (resistance), negative (susceptibility) or neutral depending on the chemical structure of the stressor. At a first attempt to understand this result, we considered the mode of action of these insecticides. The molecular targets for aldrin, deltamethrin and diazinon are the GABA gated chloride channel (Rdl), the voltage gated sodium channel and the acetylcholinesterase, respectively (Casida, 2009). Among these three proteins, only Rdl is known to interact with the PKG. The expression level of Rdl is under the control of the PKG (Kent et al., 2009) and Rdl can also be phosphorylated by the PKG (Francis et al., 2010). We have no more clues to link the toxicological features of the rover and sitter strains with the PKG and the molecular targets of the insecticides. Furthermore, the cellular locations of the PKG (intracellular) and the acetyl-choline esterase (extracellular) are not in favour of an interaction between these two proteins. So, an interaction between the PKG and these insecticides receptors is very unlikely to explain the modulation of the adaptation ability of the flies to xenobiotics.

Then, we put forward the hypothesis that the metabolization of the insecticides could explain the toxicological results. So we decided to measure activities of ESTs, GSTs and P450 enzymes families to verify our hypothesis. Since ESTs and GSTs activity levels were not able to differentiate clearly between rover from sitter flies, we concluded that they did not participate significantly to the adaptive process we study here. Contrary to these observations, the ECOD activities carried by cytochrome P450s were significantly higher in the rover strains (*for^R^* and *dg2*-cDNA) compared to their sitter counterparts (*for^S2^* and *w^1^;for^S^* respectively). Furthermore, these variations in the cytochrome P450s activities can explain the toxicological characteristics of the rover and sitter strains. Indeed, deltamethrin is inactivated by cytochrome P450s to 8-hydroxy-deltamethrin which is no longer toxic (Stevenson et al., 2011) so that rover flies should be more resistant. This assumption is verified. Aldrin is oxidized to the more toxic dieldrin by P450s (Soto & Deichmann, 1967) so that rover flies should be then more susceptible. This assumption is verified too. Finally, diazinon is metabolised by cytochrome P450s to either diazoxon (highly toxic) or diethyl-phosphate plus 2-isopropoxy-4-methyl-6-pyrimidine (not toxic) (Pisani-Borg et al., 1996) and this should result in balanced toxicological tests. This last assumption is also verified. The cytochrome P450s inhibitor PBO abolished the toxicological differences we observed between the rover and the sitter flies when in presence of aldrin or deltamethrin confirming the role of the P450s in the toxicological profile of the rover and sitter flies. Thus, the *for* gene controls the adaptation ability of flies mainly through the activity level of cytochrome P450s. It is noteworthy that the impact of this regulation is positive or negative depending on whether the xenobiotic encountered.

Because of the involvement of the cytochrome P450s *Cyp6a2* and *Cyp6g1* in metabolic resistance to insecticides in Drosophila, we traced their expression levels in the rover and sitter strains. The *Cyp6g1* gene was found over expressed in *for^R^* and *for^S2^* compared to *dg2*-cDNA and *w^1^;for^S^* but this feature does not match with neither toxicological nor P450 activities variations. We also showed that the insertion of the transposable element *Accord* fragment was present in *for^R^* and *for^S2^* at the site previously demonstrated to induce the over-expression of *Cyp6g1 (Chung et al., 2007).* From all these data, we concluded that *Cyp6g1* expression levels were independent from the PKG function and thus not involved in the adaptation ability variations we observed between rover and sitter flies. On the contrary, the *Cyp6a2* gene was found over-expressed in *for^R^* compared to *for^S2^* and in *dg2*-cDNA compared to *w^1^;for^S^* and its expression levels matched with both toxicological and activity measures results. Furthermore, we knew that the CYP6A2 protein was able to metabolize ethoxycoumarine as well as diazinon, aldrin and deltamethrin ((Dunkov et al., 1997); Amichot *et al.,* unpublished data). Although we did not demonstrate that *Cyp6a2* was an effector of the *for-*controlled adaptation to environment, we evidenced here that *for* was able to modulate the expression level of this cytochrome P450.

In the literature, references established indirect links between the PKG and the expression regulation of P450s. Previous works done with primary hepatocyte cultures from mice (Galisteo et al., 1999; Marc et al., 2000) suggested that the PKG is involved in the positive regulation of phenobarbital-induced expression of CYP2b9/10 or CYP3A P450s. A recent publication demonstrated that the PKG can regulate the function of Nrf2 (C. Chen et al., 2016). Besides, we know that the expression of *Cyp6a2* is controlled by phenobarbital (Brun et al., 1996; Sun et al., 2006) and by the Nrf2 pathway (Misra et al., 2013). All these data make sense with our present data which evidence a direct link between the PKG and the expression of the P450 *Cyp6a2*.

Interactions between environmental variations and PKG have already been documented in Drosophila. PKG plays a role in the thermotolerance (A. Chen et al., 2011; Dawson-Scully et al., 2010) and in anoxia resistance (Dawson-Scully et al., 2010). The foraging behaviour also depends on food availability (Burns et al., 2012). Interestingly, another work established a different kind of link between adaptation to the environment and the *for* gene. The authors described the identification of polymorphic regions of the genome of *Drosophila melanogaster* in relation with adaptation to the environment (Turner et al., 2008) and one of the genes identified as polymorphic in relation to environmental selection pressure (temperate vs subtropical climate) was *for*. So our studies strengthen the *for* gene as a major actor in the adaptation of drosophila to its environment because we evidence here that PKG plays a role in the adaptation to chemical stresses.

Such a relationship can have important ecological consequences. We demonstrated here that a single gene can have opposite effects on the adaptation ability of the flies depending on their foraging behaviour and on the compound they have to deal with. Because a natural population is composed of rover and sitter flies (Fitzpatrick et al., 2007; Sokolowski et al., 1997), we can suppose that a beneficial outcome should be possible for the population whatever the compound present in the environment.

A next step would be to verify whether the link we evidenced between the foraging behaviour and the adaptation ability is encountered in other insect or animal species. For instance, gregarious locusts have more PKG activity than solitary ones (Lucas et al., 2010). So this work may provide bases for future prospects about adaptation ability of migrating or invasive insects species. Molecular mechanisms linking PKG to P450 expression are also to be elucidated. Such adaptation ability in relation with foraging behaviour but not with gene induction by xenobiotics is original and deserve further studies. For instance, what could be the relevance of such a relationship between foraging and stress adaptation in insecticide resistance dissemination? Furthermore, this work focussed on chemical stress, but what about biological stress?

## Acknowledgements

We want to thank first Prof. Marla Sokolowski (University of Toronto, Canada) for providing us with the *dg2*-cDNA, *for^R^, w^1^;for^S^, for^S^* and *for^S2^* flies. We would like to specially mention Miss Alexia Lebleu for her very efficient technical support and Dr. Robichon for fruitful discussions. Finally, we thank Dr. Wajnberg for our discussions around several statistical aspects of the analysis of the experiments. This work was partly supported by a Santé des Plantes et Environnement research department (SPE, INRAE) grant (AO 2005).

## Supplementary data

**Supplemental figure 1.**
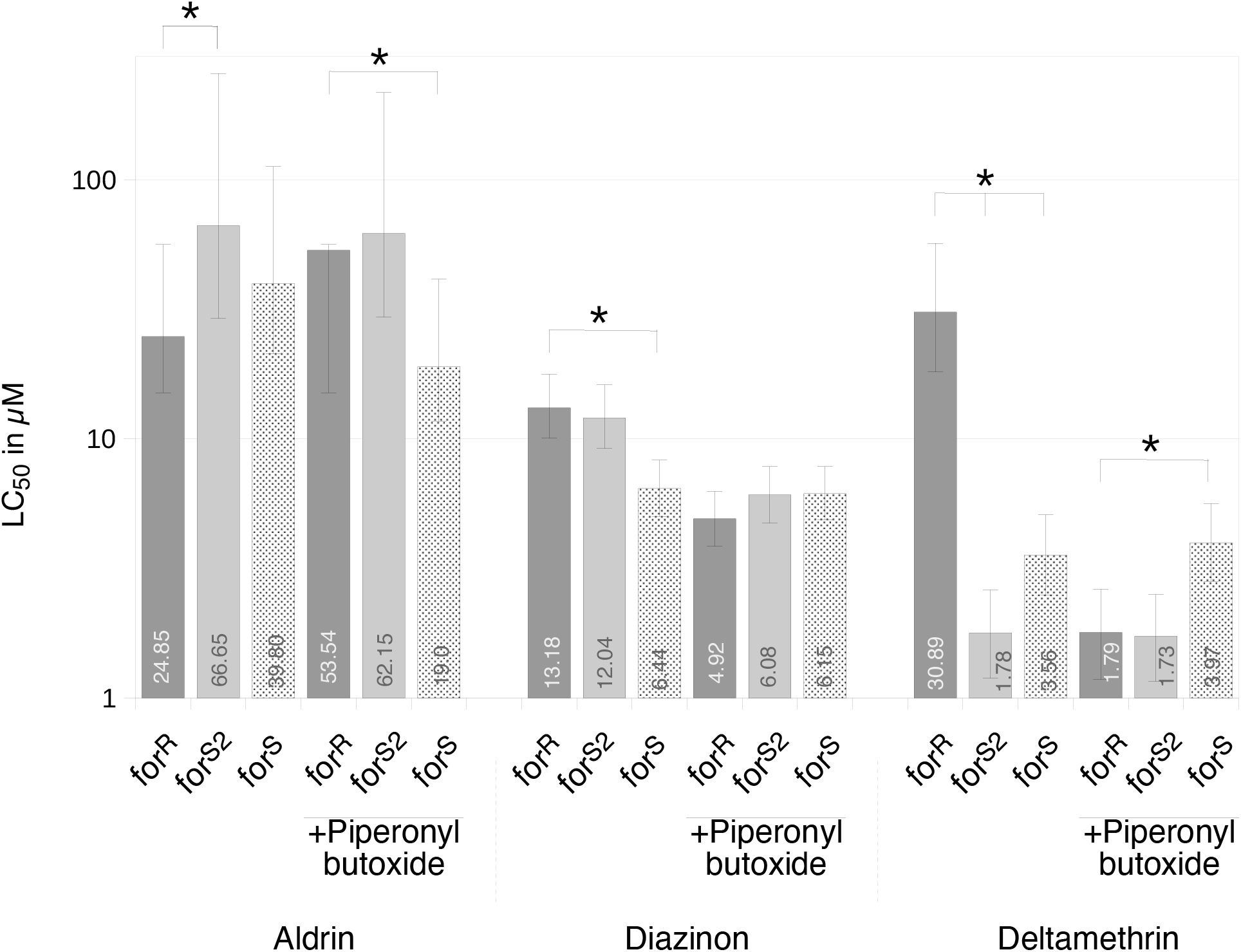
LC_50_ values of the rover and sitter strains for three insecticides Bar charts represent the LC_50_ values of forR, forS2 and forS strains. The LC_50_ values are given inside or above the boxes and are expressed in the same units as the insecticide concentrations of the acetonic solutions used to coat the tubes. The vertical bars represent the 95% fiducial limits of the LC_50_ values. The dark grey box indicate the forR strain, the light grey box the forS2 one and the dotted box the forS strain. The star indicate significantly different LC_50_ values.

**Supplemental figure 2.**
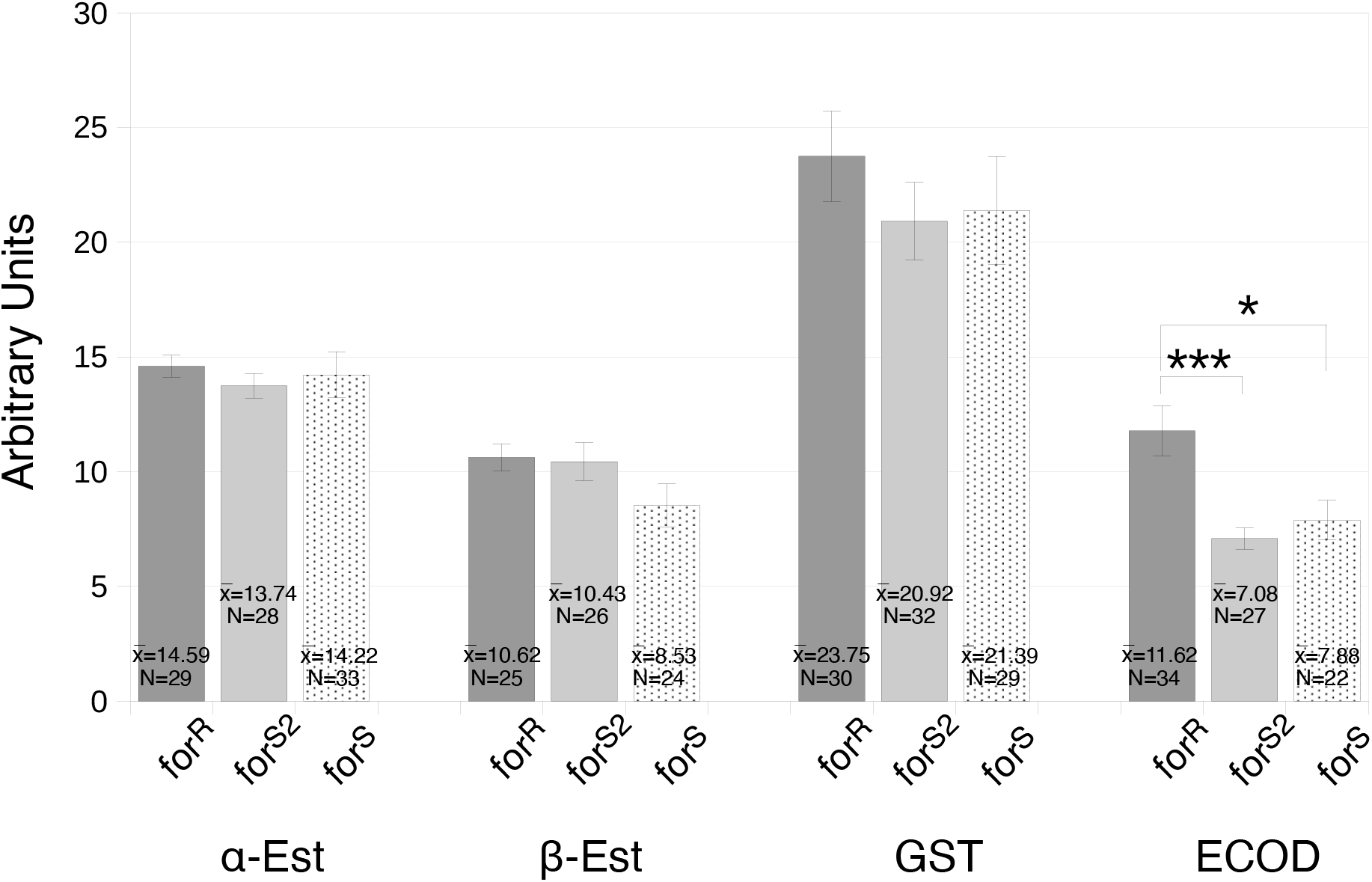
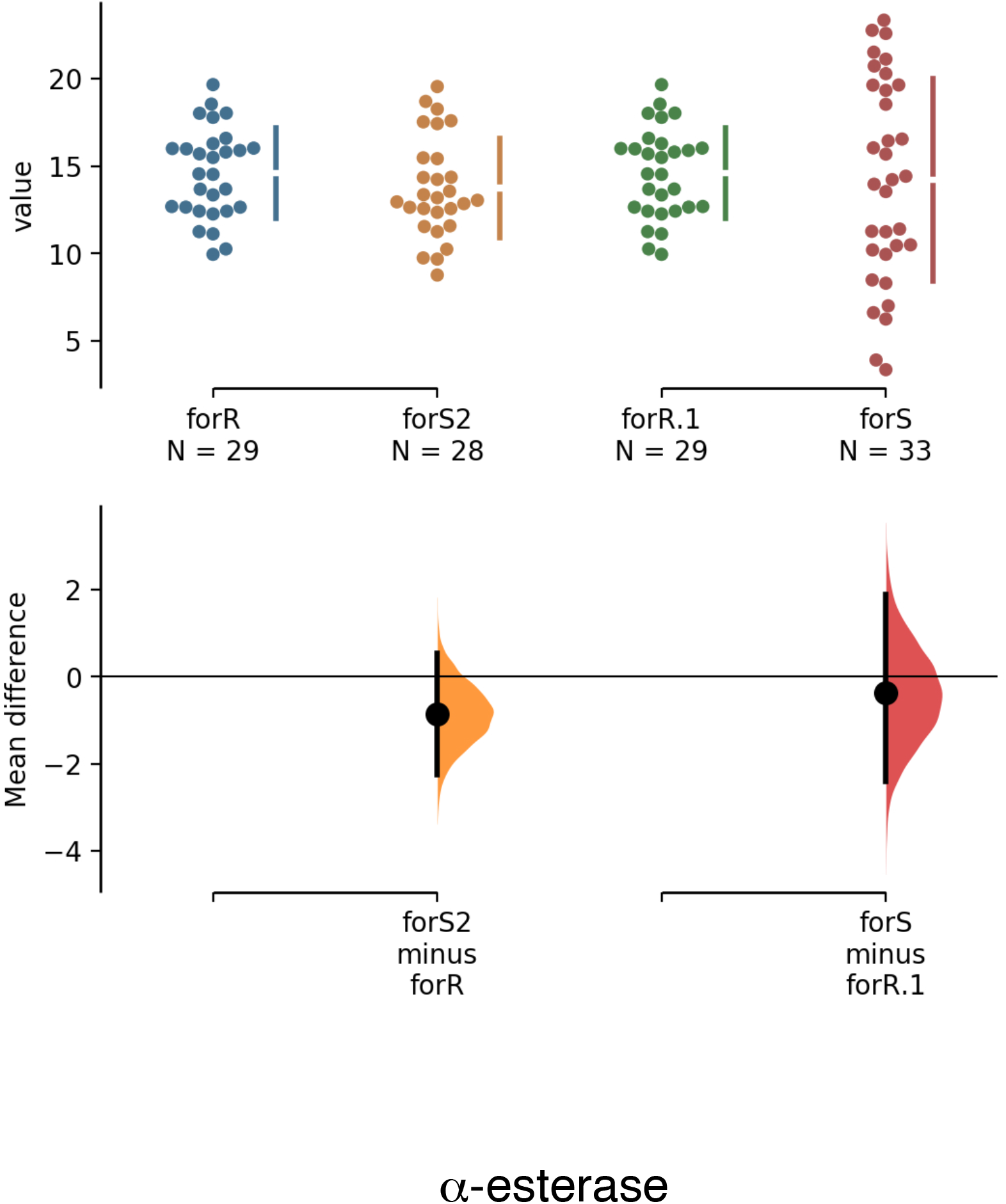
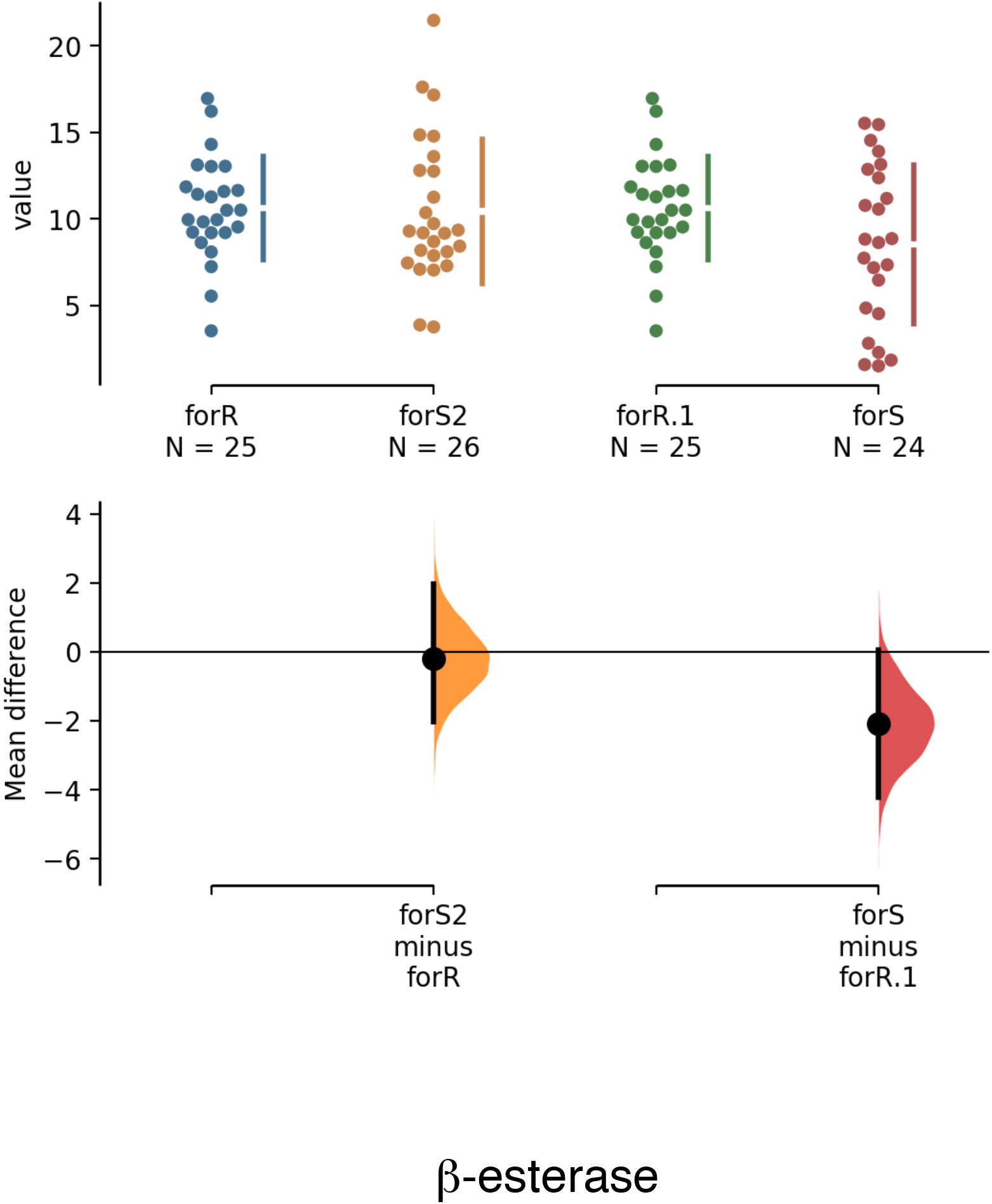
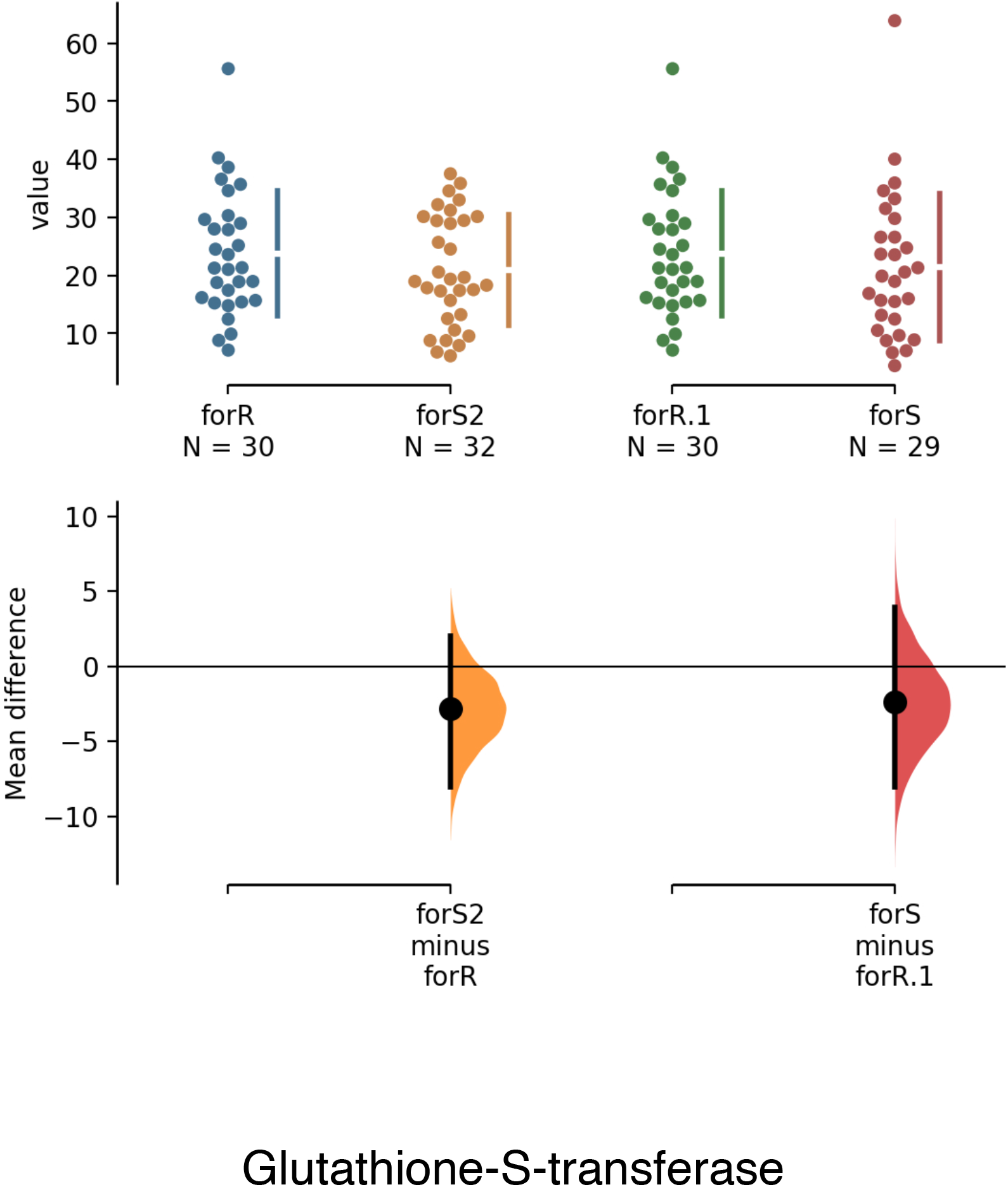
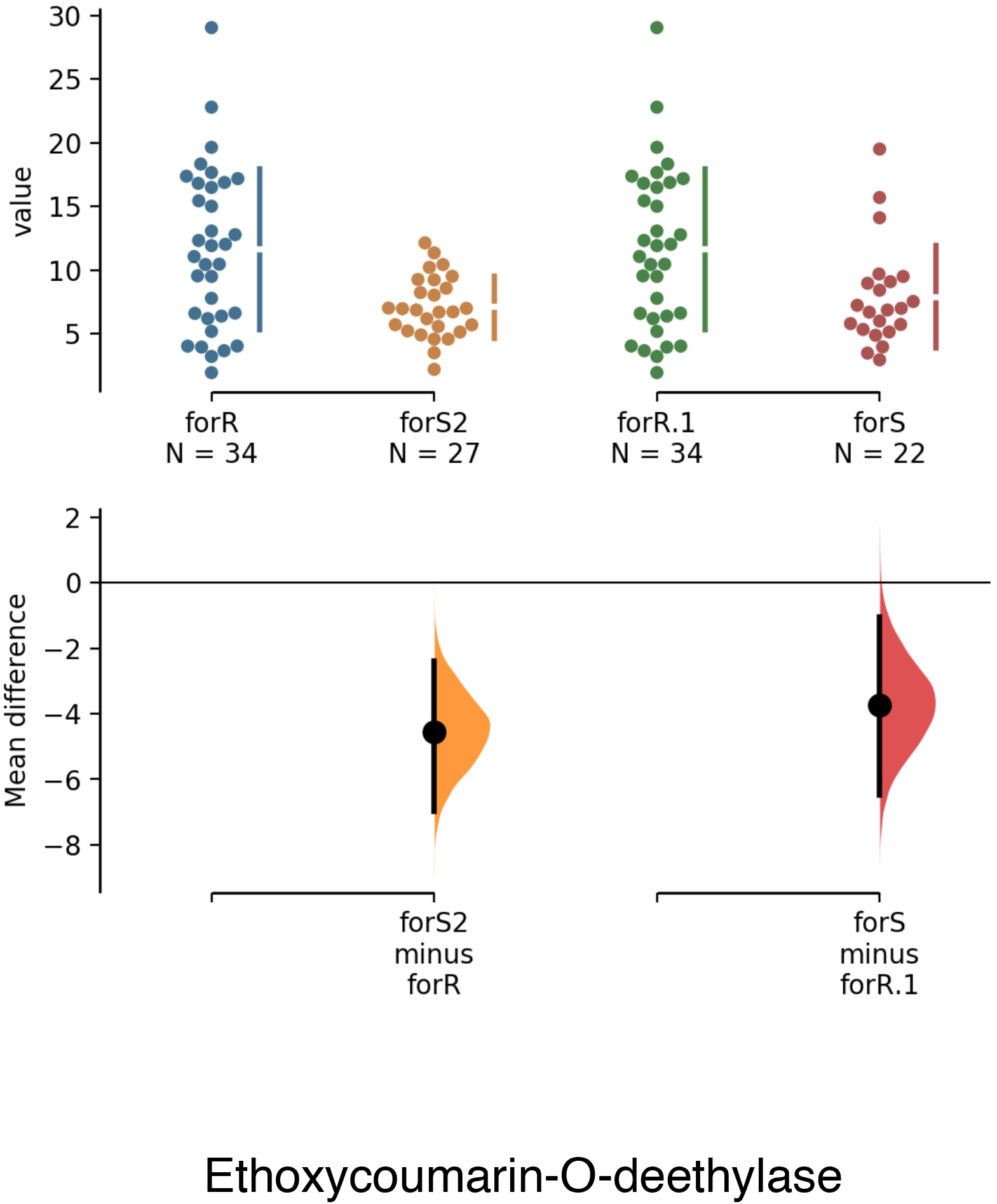
Esterases, GST and ECOD activity levels. Esterases activities are expressed as O.D. arbitrary units/fly/30 min, GST activity is expressed as fluorescence arbitrary units/abdomen/40 min and ECOD activity is expressed as fuorescence arbitrary units/abdomen/120 min. The panel A presents a bar chart of the enzymatic measurements for the forR (dark gray boxes), forS2 (light gray boxes) and forS (dotted boxes) strains. The vertical bars represent the standard error of the measures. The mean values 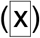 and the number of measurements (N) are given in each box. The three stars indicate the activity values significantly different at p<0,005. We also performed the Gardner-Altman two-groups estimation plot (Gardner MJ, Altman DG. 1986. Confidence intervals rather than P values: estimation rather than hypothesis testing. Br Med J. 292, 746–50) using the https://www.estimationstats.com web site facilities. The results are presented in panels B to E) for each enzymatic activity. Differences between randomly values picked in the reference set (rover strain) and in the test set (sitter strain) are calculated (5,000 rounds) and the 95% confidence interval is calculated for these differences. The presentation combines two graphs: one with the values and their means and the other with the distributions of the differences between the values after bootstrapping and the confidence interval of these distributions.

**Supplemental figure 3.**
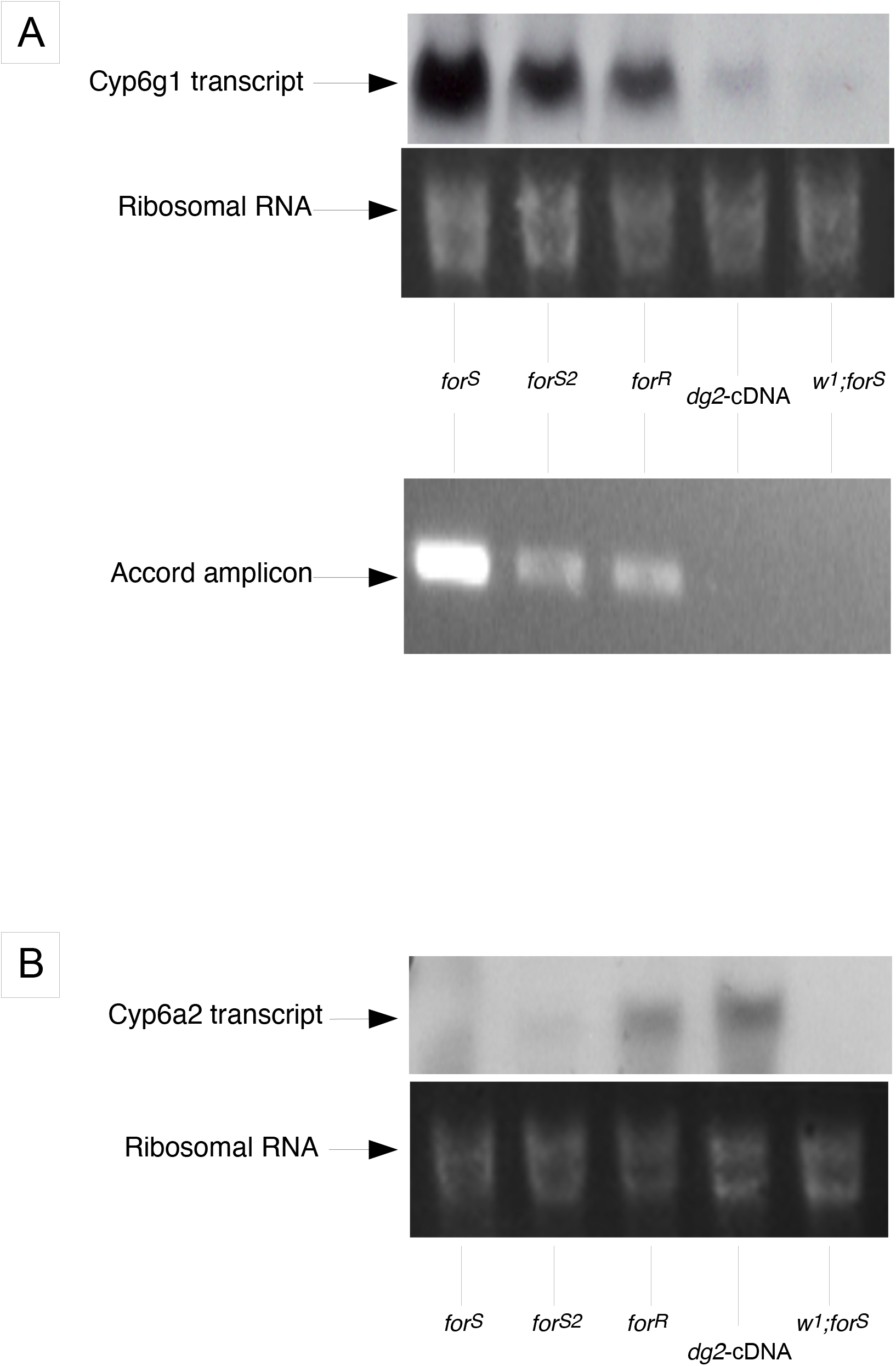
Supplemental figure 3. Expression profiles of *Cyp6g1* and *Cyp6a2* (A) Northern blot detection of the *Cyp6g1* transcript and PCR tracking of the Accord insertion in the *Cyp6g1* promoter. (B) Northern blot detection of the *Cyp6a2* transcripts. In both panels, the 18S RNA insert demonstrates a similar RNA load in the lanes.

